# T-cell potential for CD19-expressing malignancies revealed by multi-dimensional single-cell profiling

**DOI:** 10.1101/2022.06.22.497245

**Authors:** Ali Rezvan, Gabrielle Romain, Mohsen Fathi, Darren Heeke, Melisa Martinez-Paniagua, Xingyue An, Irfan N Bandey, Arash Saeedi, Fatemeh Sadeghi, Kristen Fousek, Nahum Puebla-Osorio, Laurence J.N. Cooper, Chantale Bernatchez, Harjeet Singh, Nabil Ahmed, Mike Mattie, Adrian Bot, Sattva Neelapu, Navin Varadarajan

## Abstract

Adoptive immunotherapy with T cells expressing chimeric antigen receptors (CARs) for B-cell malignancies serves as a model for identifying subsets with superior clinical activity. We profiled the infusion products (IP) of 16 patients with large B-cell lymphoma (LBCL) using an integrated suite of single-cell assays to reveal the therapeutic potential of CD19-specific CAR^+^ T cells. Timelapse imaging microscopy in nanowell grids (TIMING) profiling revealed that T cells from responders showed migration (persistent motion for at least one body length), and migration was associated with serial killing capacity. In addition, confocal microscopy revealed that migration is linearly correlated with both mitochondrial volume and lysosomal volume; and scRNA-seq demonstrated that T cells from responders were enriched in pathways related to T-cell killing, migration and actin cytoskeleton, and TCR clustering. A marker-free sorting strategy enriched T cells with migratory capacity and validated serial killing, bioenergetics, and in vivo efficacy. In aggregate, we demonstrate that migration is a cell-intrinsic biomarker independent of CAR design or biomanufacturing, desired in the bioactivity of CAR^+^ T cells associated with clinical antitumor efficacy.

## INTRODUCTION

The ability to migrate to and within tissues differentiates effective T-cell therapeutics from other targeted immunotherapies infusing recombinant proteins or small molecules. Few tools are available to examine the migratory properties of T cells despite migration being a central feature of their ability to recycle effector functions throughout malignant masses. The administration of immune effector cells propagated ex vivo has been shown to be effective for the treatment of solid tumors such as melanomas and liquid tumors such as acute and chronic B-cell leukemias^1,2^. T cells stably endowed with a genetically encoded chimeric antigen receptor (CAR) targeting CD19 have shown remarkable clinical responses in B-lineage leukemias and lymphomas patients who were refractory to other treatments. This has spurred the development of CARs targeting antigens other than CD19 to treat hematologic malignancies and invasive cancers^3-5^. The field of CAR T cells has exploded with the culmination of the Food and Drug Administration (FDA) approval of CAR T cell products, and while attention has been devoted to antigen discovery and CAR design^2,3,6^, identifying metrics that define the functional potential and thus the therapeutic prospects of T-cell products is limited^7,8^.

Because of inter- and intra-tumor heterogeneity, technologies that aggregate T-cell biology are unable to accurately capture the complexities of an IP with defined and desired characteristics. For example, populations of less differentiated cells (central memory [TCM] or stem memory [TSCM] T cells) have increased proliferative capacity leading to sustained presence, but individual cells vary in their persistence and functional potential^9-11^. Although persistence of infused T cells correlates with anti-leukemias effects, recent pre-clinical data suggest that the ability of cells to recycle effector function within the tumor microenvironment (TME) is an essential attribute for tumor eradication^12,13^. We hypothesized that functional single-cell profiling of IPs by quantifying the dynamics of T cells interacting with tumor cells would reveal properties of T cells associated with antitumor efficacy and clinical benefit.

To test this hypothesis, we performed multi-omic dynamic profiling to understand the heterogeneity of individual CD19-specific CAR T cells within axicabtagene ciloleucel (axi-cel) IP administered to patients with LBCL. By integrating function, phenotype, transcriptional profiling, and metabolism, we have identified that migration is a feature of T cells associated with clinical response and that migration is associated with bioenergetically fit cells with serial killing functionality. The link between persistent migration and killing is preserved across different CAR designs and manufacturing protocols, and provides new insights into the cellular attributes of T cells essential for efficacy.

## RESULTS

### Migratory T cells are enriched in CAR T cell infusion products (IPs) associated with clinical responses

To investigate the importance of CAR T cells characteristics, we used cells from infusion products of patients with Diffuse Large B cell Lymphoma (DLBCL) who received anti-CD19 CAR T cell therapy. In total, samples from 16 patients were collected and tested at the single-cell level using TIMING, confocal microscopy, and single-cell RNA sequencing (**Figure 1**). At 3-months follow-up after the treatment, 10 patients had a complete response (CR), and 6 showed either partial response or progressive disease (PR/PD).

**Figure 1.**
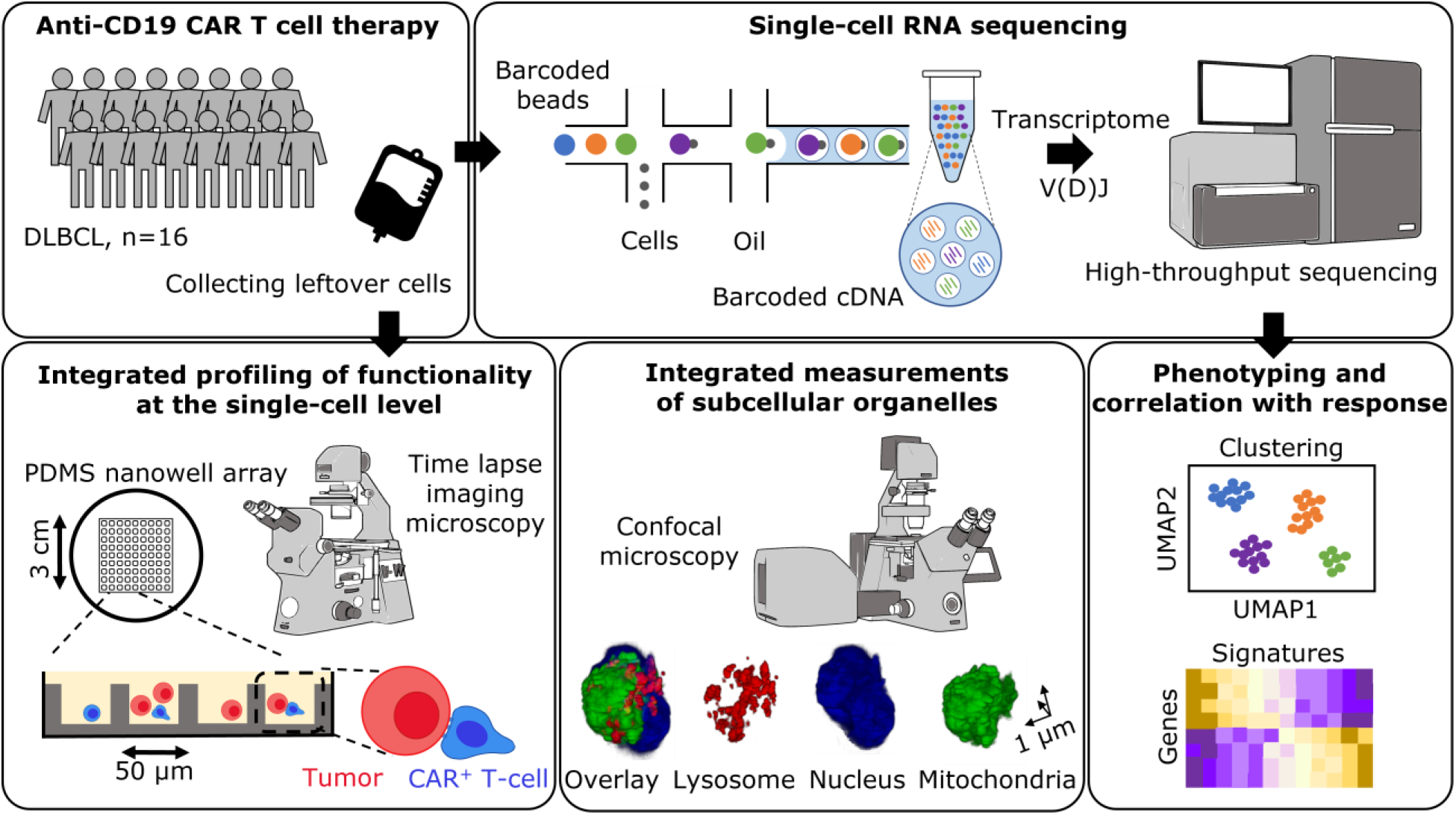
Study design for integrated single-cell multi-omic profiling of patients’ infusion products. Schematic overview of the experimental design for profiling the residual CAR T cell infusion products of 16 DLBCL patients (10 CR, 6 PR/PD). Cells were used for scRNA-seq analysis, confocal microscopy, and Timelapse Imaging Microscopy In Nanowell Grids (TIMING).

Phenotypic characterization of the CAR T cells by flow cytometry showed no differences in frequency of CAR expression [40-80%] and a balanced CD4/CD8 distribution (**Figure S1A-B**). We used the TIMING platform to dissect functional heterogeneity at the single-cell level across these 16 IPs. At an effector-to-target (E:T) of 1:1, the frequency of T cells establishing a synapse or killing the engaged tumor cells was not different between IPs associated with any clinical response (**Table ST1, Figure S2A** and **Movie M1**). At the single-cell level, the dynamics of the interaction between the T-cell and the tumor cell leading to killing were largely conserved, consistent with the expectation that intrinsic T-cell killing mechanisms are not different between the two groups of responders (**Figure S2C**). We next evaluated the ability of T cells to recycle effector (lysis) function by examining nanowells with 1E:2T and observed that the frequency of serial killer T cells was significantly enriched in IPs associated with clinical response (**Figure 2A, Figure S2B, Movie M2** and **Movie M3**).

**Figure 2.**
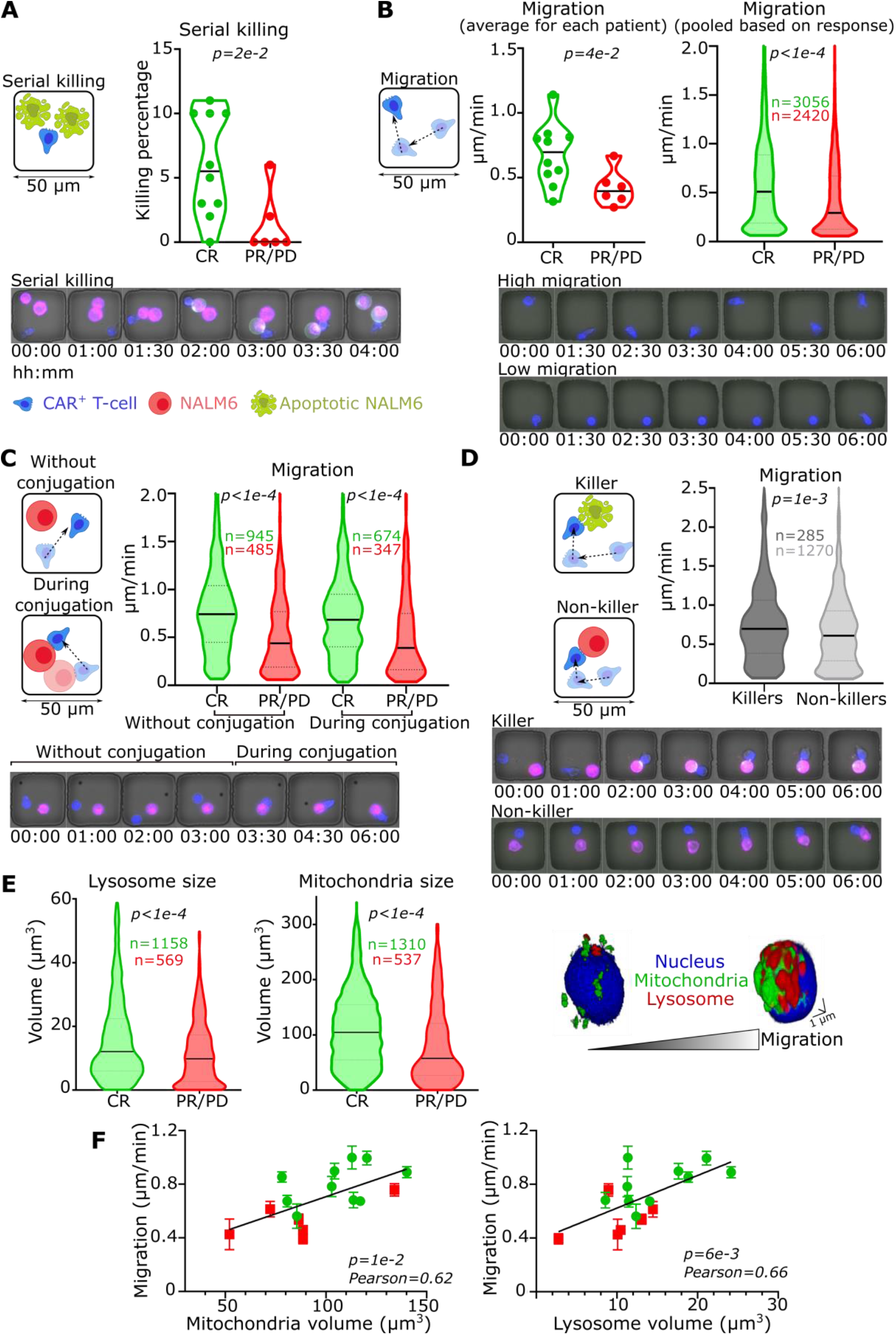
T cells from CR patients were enriched for migration, serial killing, and mitochondrial volume; in comparison to T cells from PR/PD. (A) Schematic of a serial killing event wherein a CAR T cell conjugates and kills two NALM-6 cells. The plot on the right shows the comparison between T cells from either CR or PR/PD in terms of serial killing within all 1E:2T nanowells. Micrograph showing an example of a serial killing event through the 6-hours (hh:mm) time-lapse imaging. Examples of mono-killing and no-killing events are provided in supplementary figure 1. (B) Schematic of migration for a single CAR T cell (1E:0T). The plots on the right illustrate the comparison between migration of T cells from either CR or PR/PD within all nanowells with 1E:0T. On the left plot, each dot represents the average T-cell migration for each patient, while the plot on the right shows the comparison between all T cells from CR and T cells from PR/PD. Micrograph showing examples of a T-cell with high (2 µm/min) and a low migratory capacity (0.2 µm/min). (C) Schematic of migration for a CAR T cell next to a NALM-6 cell (1E:1T) in two conditions: during the conjugation of CAR T and NALM-6 and when they are unconjugated. Plot showing the comparison between the migration of CR and PR/PD CAR T cells within all nanowells with 1E:1T without or during the conjugation. Micrograph showing an example of a CAR T cell migration before and during the conjugation with a NALM-6 cell. Not all the CAR T cells make conjugation with NALM-6. (D) Schematic of migration of a CAR T cell before conjugation with NALM-6 cell (1E:1T) in killer and non-killer CAR T cells. The plots on the right show the comparison between migration (prior to conjugation) of killer and non-killer CAR T cells within all nanowells with 1E:1T where conjugation happens (regardless of response). Micrographs showing examples of a killer CAR T cell and a non-killer CAR T cell. (E) Comparison of mitochondria and lysosome size between CR and PR/PD T cells. The confocal 3D image represents the correlation between mitochondria/lysosome size and migration of the CAR T cells. The nucleus is shown in blue, mitochondria in green, and lysosome in red. (F) Plots showing the correlation between average organelle size and average migration (1E:1T, without conjugation) of T cells. Each dot represents an IP prodcut. *P* value and Pearson correlation were calculated for the linear regression. Error bars represents SEM. The black bar represents the median, and the dotted lines denote quartiles in violin plots. *P* values were computed using Mann-Whitney tests.

T cell based drugs need to actively migrate to seek and destroy tumors. We accordingly measured the migratory potential of T cells using TIMING. In nanowells that lacked tumor cells (1E:0T), the average T-cell migration (persistent motility for at least one body-length, hereafter migration) was significantly increased in IPs associated with CR (**Figure 2B**). Aggregated by response, T cells from patients with CR showed a significantly faster migration compared to T cells from patients with PR/PD at single-cell level (**Figure 2B** and **Movie M4**). Similarly, even within 1E:1T nanowells, single T cells from patients with CR showed enhanced migration both with and without conjugation to tumor cells, compared to T cells from patients with PR/PD (**Figure 2C** and **Movie M5**). We next pooled T cells from all IPs (1E:1T nanowells, regardless of clinical response) and this comparison confirmed that migration is an intrinsic feature of killer T cells compared with non-killer T cells (**Figure 2D** and **Movie M6**).

In vivo proliferative capacity leading to cellular persistence after infusion is one of known correlates of CAR T cells^14^. We integrated TIMING with 3D confocal microscopy to interrogate the correlation between migration, killing, and proliferative capacity within IP T cells. We constructed the image of each cell as a series of stacks in 3D to capture the mitochondrial volume (known feature of proliferative capacity)^14^, lysosomal volume (killing capacity), and the nucleus (cell size). When the T cells from the IPs were stratified by clinical response, T cells from CR patients had increased mitochondrial and lysosomal volume in comparison with T cells from PR/PD patients (**Figure 2E**). Since IP T cells from each patient had been characterized by both TIMING and confocal microscopy, we investigated the correlation between migration, mitochondrial and lysosomal volumes. We observed a strong linear correlation between migration and mitochondrial mass (Pearson correlation 0.62, *P* value = 0.01), and between migration and lysosomal volume (Pearson correlation 0.66, *P* value = 0.006) (**Figure 2F**). We performed unsupervised hierarchical clustering of all the dynamic parameters from TIMING, and the organelle measurements from confocal microscopy. Clustering showed that serial killing, migration, mitochondrial volume and lysosomal volume were features associated with T cells from patients who achieved CR (**Figure S2D**). Collectively, these single-cell measurements illustrate that IP T cells from patients with CR, balance migration, serial killing and cellular fitness.

### ScRNA-seq identifies a core signature of migration in IP T cells from patients who achieved CR

To understand the molecular basis of these observations, we performed scRNA-seq of 21,469 T cells from nine of these patients (5 CR, 4 PD) for whom we had access to additional T cells from the IPs. To minimize batch effects, we barcoded samples in sets of three, prepared libraries and sequenced them together. We defined ten clusters of T cells (T1-10) based on their molecular properties and cells derived from patients with both CR and PD were represented in all the clusters (**Figure 3A**). Consistent with previous reports, the CAR transcript was distributed in all of these different clusters, but the overall frequency of the detected transcript was lower than CAR expression by flow cytometry, suggesting a high rate of dropout of the CAR transcript (**Figure S1A**)^8^. Accordingly, we performed subsequent analyses on all T cells from IPs.

**Figure 3.**
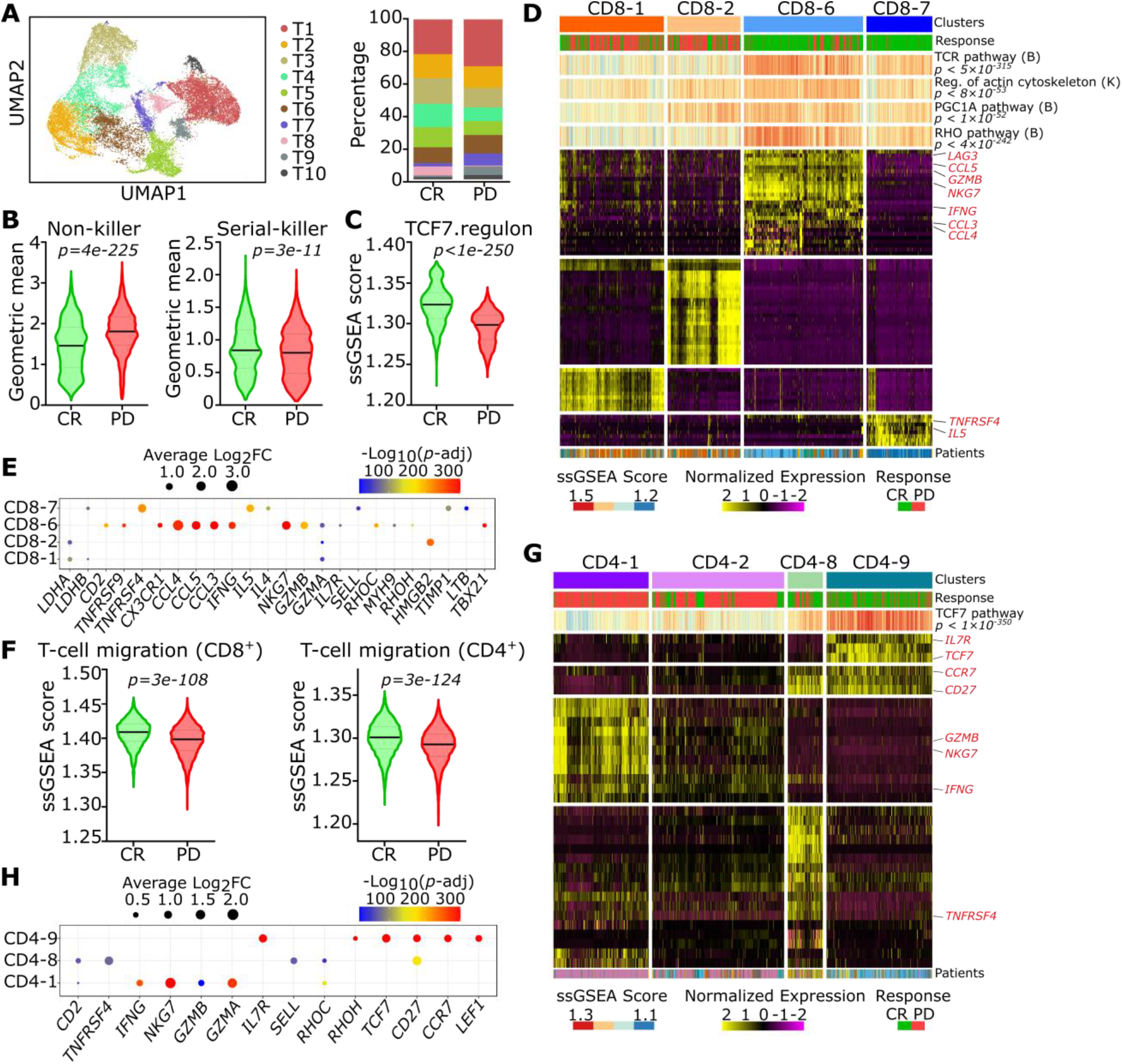
Transcriptional profiling of CAR T infusion products associated with response revealed by single-cell RNA sequencing. (A) Uniform Manifold Approximation and Projection (UMAP) for 21,469 cells from nine IPs. Bar plot showing the distribution of T cells from CR and PD/PR among 10 clusters determined using unsupervised clustering. (B) Comparison between CD8^+^ T cells from CR and PD for non-killer (*GZMA* and *CD69*) signatures or serial-killer (*CD2* and *CD27*) T cells. (C) Comparison between CD8^+^ T cells from CR and PD for TCF7 regulon signatures calculated using ssGSEA. (D) Heat map of four CD8^+^ T cell clusters generated by unsupervised clustering. CD8-1 and CD8-2 contain mostly cells from PD while CD8-6 and CD8-7 are enriched with CR cells. A color-coded track on top shows the cells from infusion products of CR (green) and PD (red). Additional tracks show the ssGSEA scores of TCR, actin cytoskeleton regulation, PGC1A, and RHO pathways, respectively. B: BIOCARTA and K: KEGG show the source for the pathways. These pathways are significantly enriched in CR clusters compared to PD clusters. The track below the heatmap shows the sample origin for each cell. (E) Bubble plot showing key genes differentially expressed among four CD8^+^ T clusters. (F) Comparing ssGSEA-derived migration score between T cells from CR and PD for either CD4^+^ T cells or CD8^+^ T cells. (G) Heat map of four CD4^+^ T cell clusters generated by unsupervised clustering. CD4-1 and CD4-2 contain mostly cells from PD while CD4-8 and CD4-9 are enriched with CR cells. A color-coded track on top shows the cells from infusion products of CR (green) and PD (red). Additional track shows the ssGSEA scores for TCF7 regulon pathway which is significantly enriched in CR clusters compared to PD clusters. The track below the heatmap, shows the sample origin for each cell. (H) Bubble plot showing key genes differentially expressed among CD4^+^ T clusters. For violin plots, the black bar represents the median and the dotted lines denote quartiles. *P* values for both heatmaps and violin plots were computed using Wilcoxon tests.

We did not observe a significant expression of either well-established genes associated with T-cell exhaustion including *TOX, PDCD1, TIGIT*, and *CD38*; or emerging regulators like *ID3* and *SOX4*, within any of our T cells clusters (**Figure S3A**)^8,15^. We next focused the analysis on CD8^+^ T cells. Using multi-omic dynamic profiling of healthy donor-derived CD19-specific CAR T cells, we have recently published a molecular signature of both non-killer and persistent serial killer CD8^+^ T cells^16^. We performed single-cell gene set enrichment analysis (ssGSEA) and showed that consistent with our functional data (**Figure 2A**), serial killer T cells were significantly enriched in IPs from patients with CR whereas non-killers were significantly enriched in IPs from patients with PD (**Figure 3B**). CD8^+^ T cells from patients who achieved CR also showed enrichment in a *TCF7* gene signature (**Table ST2**) associated with enhanced in vivo persistence compared to CD8^+^ T cells from patients with PD (**Figure 3C**)^17^. We next identified differentially expressed genes (DEGs) between CD8^+^ T cells from IPs that resulted in CR compared to IPs of patients with PD. We performed unsupervised clustering based on the DEGs and identified seven clusters (**Figure S3B**). Clusters CD8_1 and CD8_2 were predominantly comprised of cells from PD, whereas clusters CD8_6 and CD8_7 were predominantly comprised of cells from CR (**Figure 3D**). Cells within the CD8_6 cluster showed a high expression of genes associated with effector functionality including cytotoxicity (*GZMB, PRF1, FASLG*, and *NKG7*), cytokine and chemokines (*CCL3-5, IFNG*), and migration (*MYH9, RHOH*, and *RHOC*) (**Figure 3E**). Cells in clusters CD8_6 and CD8_7 were significantly enriched in pathways associated with TCR activation (*p* < 5×10^−315^), PGC1A and mitochondrial biogenesis (*p* < 1×10^−52^), actin cytoskeleton regulation (*p* < 8×10^−53^) and migration (RHO pathway, *p* < 4×10^−242^) (**Figure 3D**). This overlap between TCR activation, cytotoxicity and migration at the single-cell level is consistent with the known immunobiology of T cells, the same genes that control cellular migration also control cytoskeletal remodeling at the immunological synapse.

To map specificity to T cell migration, we curated a collection of 40 genes (labeled T-cell migration score, **Table ST2**) with documented roles in actin cytoskeleton remodeling in T cells^18^ including *RHOA* and *MYH9* (actomyosin contraction), *TLN1* and *VCN* (actin-integrin interplay), and *MYO1G* (actin contraction). ssGSEA confirmed that the T-cell migration score was significantly enriched in the CD8_6 and CD8_7 clusters compared to the CD8_1 and CD8_2 clusters (*p* < 5×10^− 146^) (**Figure S3B**). CD8^+^ T cells from patients who achieved CR showed a significant enrichment in the T-cell migration score compared to CD8^+^ T cells from patients who associated with PD (**Figure 3F**). It has been recently reported that T-cell migration signatures can also be associated with T-cell exhaustion^19^. Within our dataset, the T cells, regardless of response status, did not show significant expression of the genes associated with the migration/exhaustion phenotype including *CD38, MYO7A, MYO7B*, or *CAV1* (**Figure S3A**).

We next analyzed CD4^+^ T cells and first confirmed that CD4^+^ T cells from patients who achieved CR showed a significant enrichment in the T-cell migration score compared to CD4^+^ T cells from patients who associated with PD (**Figure 3F**). Analysis of the DEGs and unsupervised clustering of the CD4^+^ T cells revealed nine clusters (**Figure S3C**). Two clusters, CD4_1 and CD4_2, predominantly comprised cells from PD, whereas clusters CD4_8 and CD4_9 were predominantly comprised of cells from CR (**Figure 3G**). In contrast to the CD8^+^ T cell clusters, genes for immediate cytotoxicity were not high in CD4_8 and CD4_9; rather these clusters showed an enrichment of genes associated with long-term in vivo persistence including *LEF1, TCF7, CD27*, and *IL7R* (**Figure 3H**). In aggregate, scRNA-seq revealed that CD8^+^ T cells from IPs with CR are enriched in serial killers and associated with immediate cytotoxicity whereas CD4^+^ T cells from IPs with CR are associated with long-term persistence. This divergence in function and long-term in vivo persistence is well-supported by other pre-clinical and recent long-term follow-up clinical data^20,21^. Importantly, scRNA-seq revealed that migration is an intrinsic property of both CD4^+^ and CD8^+^ T cells associated with clinical responses.

### T-cell migration enables selection of functional T cells

Based on our results of profiling clinical IPs, we next wanted to identify if migration can serve as a biomarker to identify the fittest cells for adoptive immunotherapy. We sought proof-of-concept studies based on defined populations of healthy donor-derived CD19-specific CAR T cells to utilize migration as a selectable property for T-cell bioactivity. We hypothesized that T cells with migration could be enriched using a modified transwell assay, with pore sizes consistent with previously published confinement studies of T cells in vitro^22^. We initially tested the effect of functionalization of the membrane by coating it with fibronectin or collagen, but in comparison with the uncoated membrane, these did not significantly alter the number of migrated (migratory) cells (not shown).

Unstimulated CD19-specific CAR^+^ human T cells with CD28 endodomain (designated 19-28z), comprised of both CD4^+^ and CD8^+^ T cells, were seeded onto a Boyden chamber, and migratory T cells were harvested from the bottom chamber and compared to cells from the top chamber (non-migratory) or unsorted cells (**Figures S4** and **S5**). TIMING confirmed that the cells harvested from the bottom chamber had an increased frequency of individual T cells with migration in comparison to the cells harvested from the top chamber, confirming that at least for proof-of-concept studies the modified transwell assay could be utilized to enrich T cell populations based on the migration (**Figure 4A**). To dissect the cytotoxic capacity at the single-cell level, we performed TIMING assays with migratory or non-migratory 19-28z T cells as the effector cells and NALM-6 cells as targets. Consistent with the data from the clinical IPs, migratory T cells participated in killing and serial killing at significantly higher frequencies in comparison to the non-migratory population (**Figure 4B**). Since we recognized that the TIMING data is gated based on T cells that conjugate to tumor cells (*i*.*e*. ones that establish a synapse), and posited that migration is likely to impact the ability of T cells to seek and conjugate to tumor cells, we modeled the time evolution of the interaction and outcomes of the interaction between individual T cells and tumor cells using a state transition diagram. Consistent with our hypothesis, the most probable path for migratory T cells was to encounter multiple tumor cells and behave as serial killers, whereas non-migratory T cells were characterized by an impediment in conjugating to tumor cells (**Figure 4C**). Collectively, these results suggest that the migratory T cells isolated by the transwell assay were enriched in serial killers.

**Figure 4.**
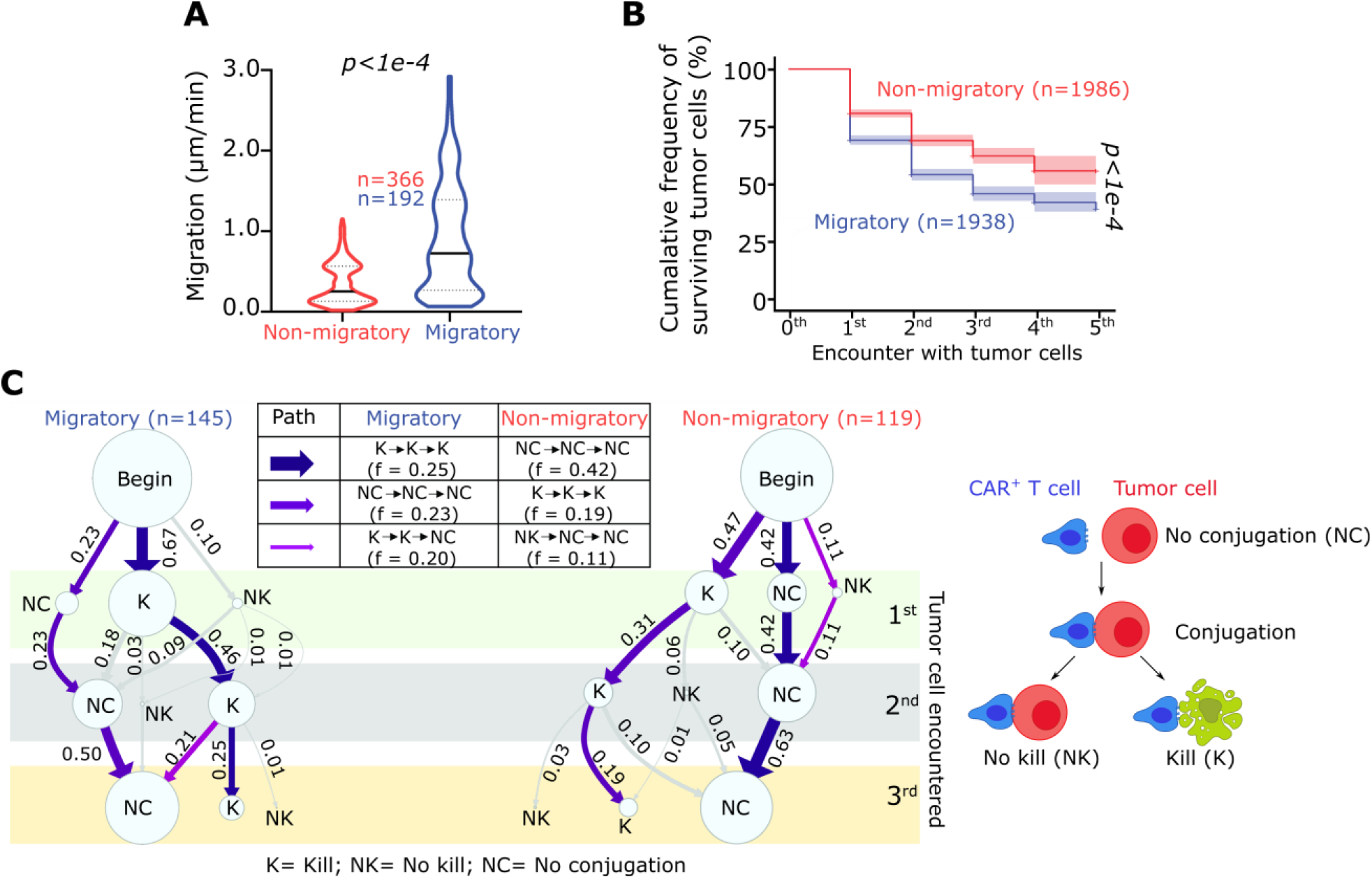
Enrichment and functional characterization of migratory 19-28z T cells. (A) Comparisons between the migration of migrated (migratory) and non-migrated (non-migratory) cells. The black bar represents the median and the dotted lines denote quartiles. *P* value were computed using Mann-Whitney tests. (B) Sustained killing mediated by individual migratory 19-28z T cells ordered by the encounter with tumor cells. Error bars indicate 95 % CI. *P* value was computed using log-rank test. (C) State transition diagram illustrating the evolution of the interaction between 19-28z T cells and tumor cells within single-cell assays. The thickness of the lines connecting the state is proportional to the frequency of the transition. The data was obtained from nanowells containing exactly one T cell and 3-5 tumor cells. All *P* values were computed using Mann-Whitney tests and each dot represents a single effector cell. All data shown here are from one representative population derived from at least three independent healthy donor-derived 19-28z T cells.

### Migratory T cells have a high spare respiratory capacity (SRC) but are phenotypically similar to the unsorted T cell populations

We next sought to determine if the sorted migratory 19-28z T cells, marked the subpopulation with superior efficacy, and to define the properties that distinguished the migratory population from the parent, unsorted population. To test if the differences in the migration and functional properties of migratory T cells could be explained by their phenotype, we compared the memory phenotype of migratory and unsorted T cells, and observed that both populations were comprised predominantly of naïve (CD62L^+^CD45RA^+^) and central memory (CD62L^neg^CD45RA^+^) CAR^+^ T cells (**Figure 5A** and **Figure S5**). Both migratory and unsorted populations had no difference in expression of either the CAR or intracellular Granzyme B (**Figure S5** and **Figure 5B**), consistent with transcriptional and functional data from IP profiling.

**Figure 5.**
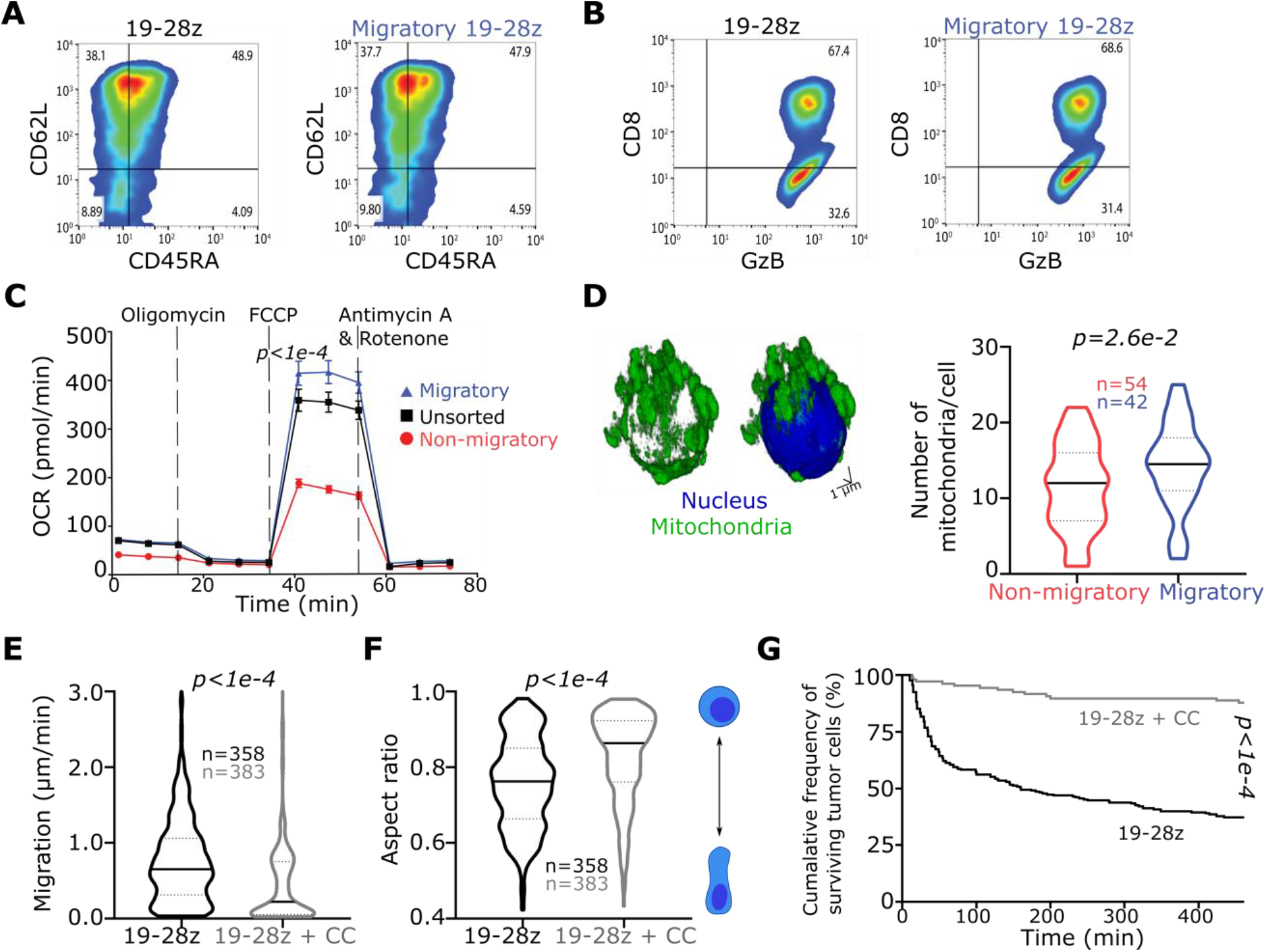
The phenotype and bioenergetics of migratory CAR T cells. (A/B) The phenotype (A) and Granzyme B (GzB) expression (B) of the migratory and unsorted 19-28z T cells as determined by flow cytometry. (C) Basal OCR levels measured for three different 19-28z T cell populations. *P* value is for comparison of the SRC comparing the migratory and non-migratory subsets using multiple t tests. (D) A confocal 3D image of a migratory 19-28z T cell. Nuclei are shown in blue and mitochondria in green. The plot shows the number of mitochondria per cell compared between migratory and non-migratory 19-28z T cells. (E/F) The migration and polarization of 19-28z T cells treated with Compound C (CC). All data representative of at least three independent experiments performed with cells from at least three healthy human donors-derived 19-28z T cells. The black bar represents the median and the dotted lines denote quartiles. The *P* value was computed using a Mann-Whitney test. (G) Comparisons of the killing frequency of vehicle treated (DMSO) or CC treated 19-28z CAR T cells. Each data point represents a single-cell. *P* value was computed using log-rank test.

Migration is an energy intensive process and the data from profiling the IPs suggested that migration in serial killer T cells is also correlated with increased metabolic fitness. To directly establish a link between migration and metabolism, we measure the oxygen consumption rate (OCR) of both the migratory and unsorted T-cell populations. Metabolic flux analyses revealed that the migratory populations had both higher maximal respiratory capacity and spare respiratory capacity (SRC) in comparison with either the unsorted or non-migrated T-cell populations (**Figure 5C**). Since the data from IPs suggested differences in mitochondrial volume, we utilized 3D single-cell confocal microscopy to study mitochondrial structure. Imaging of single cells confirmed that migratory 19-28z T cells had increased mitochondrial volume and an increased number of punctate mitochondria (consistent with fission) in comparison to the non-migratory 19-28z T cells (**Figure 5D**). Since AMP-kinase (AMPK) is a well-known regulator of mitochondrial mass and integrity, we next sought to determine if the kinase activity of activated AMPK (phosphorylation in the α-subunit) was the molecular link between migration and metabolism that we have documented. Accordingly, we inhibited the activity of AMPK in 19-28z T cells using the small molecule inhibitor dorsomorphin (compound C, CC). Cells treated with CC showed profound defects in morphology and migration, confirming that AMPK activity is essential for the migration of T cells (**Figure 5E-F**). Consistent with impaired migration, CC treated 19-28z T cells showed a decreased propensity to conjugate to tumor cells in single cell assays, in comparison to DMSO-treated 19-28z T cells (**Figure S6A**). Furthermore, treated 19-28z T cells that did make contact with target cells showed significantly longer conjugation times and delayed induction of tumor cell apoptosis when compared to the untreated 19-28z T cells (**Figure S6B** and **Figure 5G**). We also confirmed that this requirement of AMPK for migration was generalizable to genetically unmodified, tumor-reactive T cells and not just CAR^+^ T cells (**Figure S7** and **Movie M7**). Collectively, these results show that transwell-sorted 19-28z T cells phenocopy the functional properties of T cells from patients who achieved CR: migration, serial killing, and mitochondrial fitness.

### Migratory T cells reject established leukemia and sustain persistence in vivo

Based on the in vitro functional and bioenergetics data, we hypothesized that migratory T cells should be able to promote CAR^+^ T-cell persistence in vivo leading to improved antitumor efficacy. We assessed the efficacy of the migratory 19-28z T cell population using a model of established leukemia. NOD.Cg-Prkdcscid Il2rgtm1Wjl/SzJ (NSG) mice were injected with CD19^+^ NALM-6 transduced with firefly luciferase (ffLuc), and after confirming the engraftment of tumor, we treated the mice with 19-28z CAR^+^ T cells (**Figure S8**). The migratory 19-28z T cells demonstrated potent and superior antitumor activity, reducing tumor burden to the detection limit, with tumor flux significantly lower in comparison to the unsorted 19-28z T cells (**Figure 6A-B**). It is worth emphasizing that this improvement in efficacy was obtained purely by isolation of the subpopulation of migratory cells in the population without any additional culturing or modifications. In both the bone marrow (BM) and spleen, mice treated with migratory 19-28z T cells harbored no tumor cells, but only persisting CAR^+^ T cells (**Figure 6C**). By contrast, we could detect the outgrowth of tumor cells within the BM of mice treated with unsorted 19-28z T cells, which was accompanied by a low frequency of CAR^+^ T cells in the BM in these same mice (**Figure 6C**). Similarly, while we could detect CAR^+^ T cell persistence in the spleen of mice infused with migratory 19-28z T cells, there were very low frequencies of CAR^+^ T cells in the spleen of the mice that received the unsorted 19-28z T cells (**Figure 6C**). In an independent experiment, we compared the efficacy of migratory and unsorted 19-28z T cells at suboptimal doses against these same NALM-6 tumors in vivo; migratory 19-28z T cells showed enhanced antitumor activity compared to the unsorted 19-28z T cells (**Figure S9**). In aggregate, these results demonstrate that migration is a desired biomarker indicative of subpopulations of T cells with desired function and in vivo persistence.

**Figure 6.**
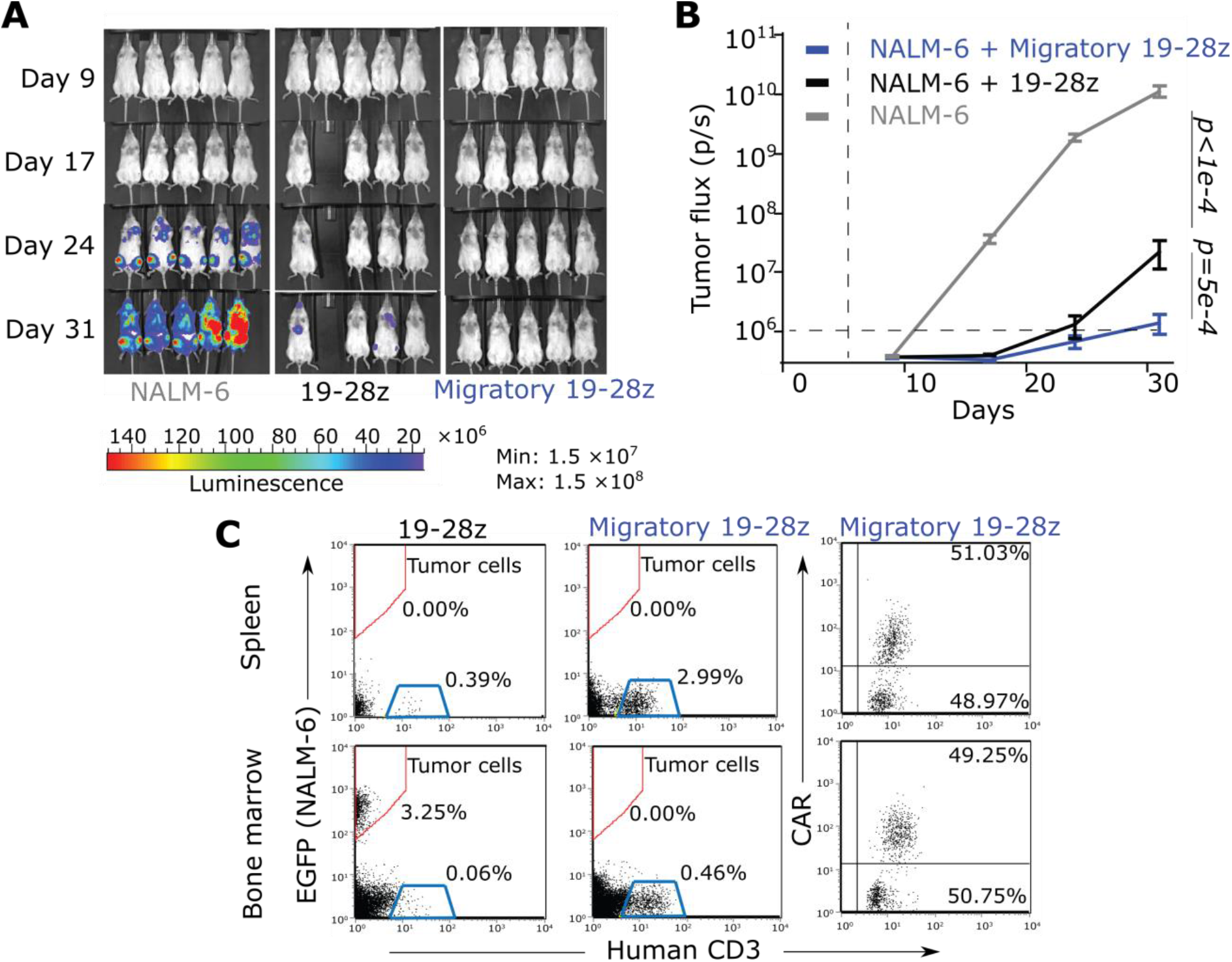
Migratory 19-28 T cells reject established leukemia and sustain persistence in vivo. (A) False-colored images illustrating the photon flux from ffLuc expressing EGFP^+^NALM-6 cells. (B) Time course of the longitudinal measurements of NALM-6 derived photon flux from the three separate cohorts of mice (n= 10 in each group). The background luminescence was defined based on mice with no tumor. Error bars represent SEM and *P* values are computed using the Mann-Whitney test. (C) On day 31, four mice from each group were euthanized, and tissues (bone marrow and spleen) were harvested and analyzed by flow cytometry for expression of human CD3 (human T cells) and EGFP (gated within hCD19 cells). The CAR^+^ T cells were identified by a scFv-specific antibody, as described previously^54^. The flow data is representative from one mouse in each group.

### Migration is a biomarker of functional T cells independent of CAR design or biomanufacturing protocols

We next tested if the link between migration and functionality is generalizable to CD19-specific CARs derived from diverse manufacturing protocols, and against multiple tumor targets. Accordingly, we utilized TIMING to first compare the migration of killer and non-killer CAR^+^ T cells electroporated and manufactured with a CD19-specific CAR containing the CD8 hinge and transmembrane regions (19-8-28z) (**Figure 7A**). Consistent with our other data, individual killer 19-8-28z T cells demonstrated higher migration both with and without conjugation to NALM-6 tumor cells compared with non-killer 19-8-28z T cells (**Figure 7B** and **S10A**). Next, we tested two tri-specific CAR^+^ T cells designed for mitigating CD19 escape in primary ALL (**Figure 7C** and **7E**)^23^. These tri-specific CAR^+^ T cells were retrovirally transduced, the CAR design incorporated a 41BB endodomain, and the T cells were expanded (anti-CD3/CD28 and IL7/IL5), as described previously^24^. We tested these T cells against patient-derived tumor cells. Again, by comparisons using TIMING, individual killer CAR^+^ T cells demonstrated higher migration both with and without conjugation to patient-derived tumor cell lines compared with non-killer CAR^+^ T cells (**Figure 7D, 7F** and **Figure S10B-C**). These data suggest that the link between migration and killing can be preserved across multiple CAR designs and manufacturing protocols.

**Figure 7.**
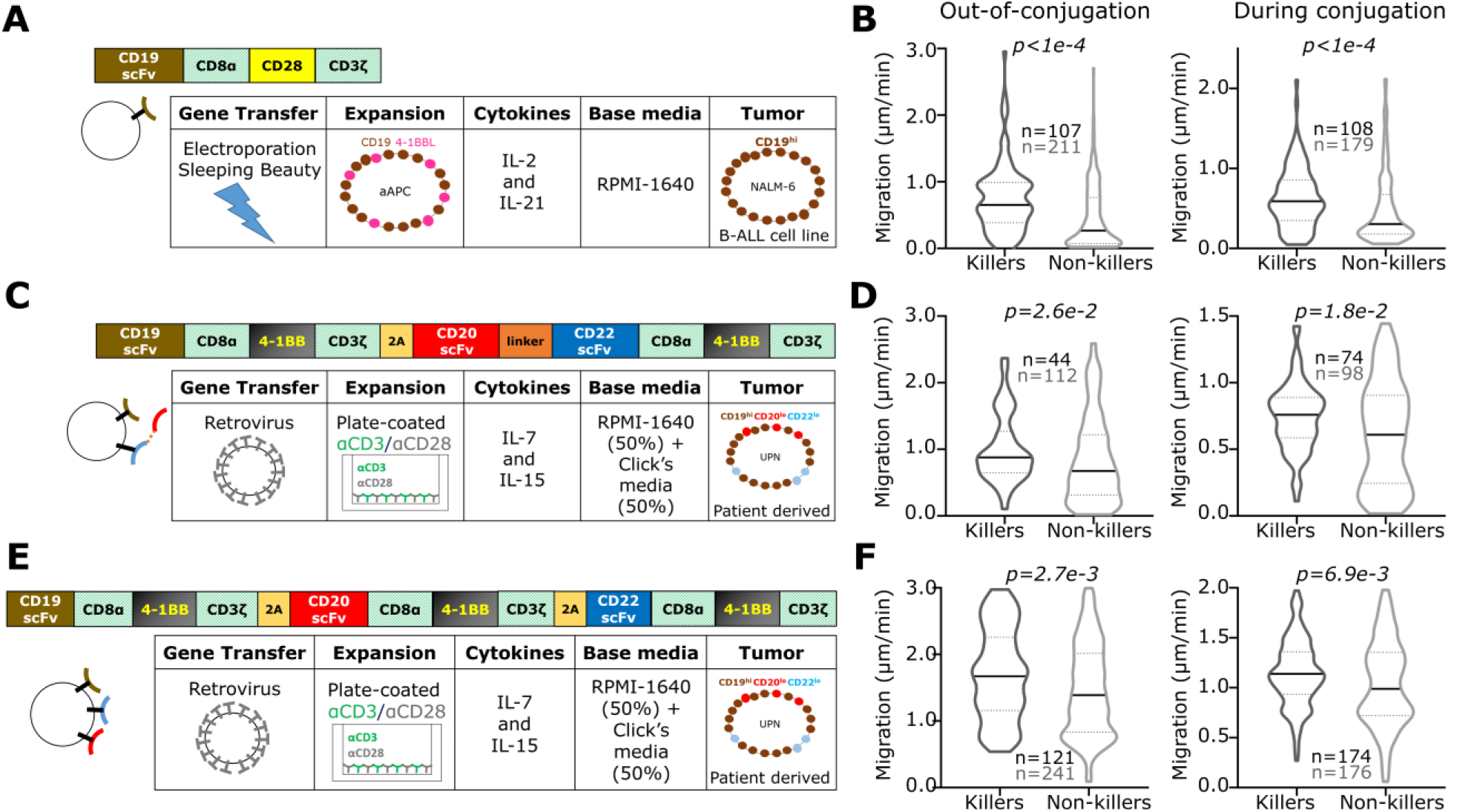
Quantifying the link between migration and functionality in diverse CARs. (A, C and E) Schematic illustrating the CAR structure, manufacturing and expansion, and the target cells used for profiling functionality of individual CAR^+^ T cells using TIMING. (B, D and F) The migration of individual killer and non-killer CAR T cells without and with conjugation to tumor cells. All data from an E:T of 1:1. The black bar represents the median and the dotted lines denote quartiles. All *P* values were computed using Mann-Whitney tests and each data point represents a single effector cell.

## DISCUSSION

With the approval of CD19-specific CAR^+^ T cells as living drugs, there is a need to define biomarkers that provide insights into the clinical impact of adoptive immunotherapy. Identifying cellular properties associated with clinical efficacy are important for ensuring predictable clinical outcomes and for defining the desirable attributes during biomanufacturing. Our results demonstrate that the variability in IP can directly impact clinical efficacy and that migration feature can be used to identify serial killer T cells with enhanced in vivo persistence. Prior studies of IPs have been limited to static profiling based on flow cytometry^25^, whole-transcriptome profiling^8,25^, or cytokine secretion^7^, but our dataset is the first multi-omic profile that includes dynamic imaging across thousands of T cells and their interactions with tumor cells. Our discovery of migration as a cellular biomarker of T cells with in vivo persistence is consistent with pre-clinical studies using two-photon microscopy that demonstrated: (1) mouse T-cell migration prior to engagement and killing of tumor cells is an essential component of their efficacy^26,27^, (2) both CD4^+^ and CD8^+^ CAR^+^ T cells mediate direct and indirect cytotoxicity upon migration to the tumors^28^, (3) migratory T cells are long-duration tumor-resident T cells^19^, (4) functional exhaustion by PD-1 induces T cell migration paralysis and blockade of PD-1 can restore T cell migration and function^29^, and (5) T-cell migration in tissue increases with serial killing and tumor rejection^30^. This ability of intrinsic T cell migration to promote tumor exploration is a central feature of cells as drugs that distinguishes them from small molecule and protein therapeutics^31^. At the molecular level, the genes/proteins involved in migration, including *RHOA, RAC1*, and *CDC42*, affect multiple aspects of T-cell immunobiology, including cell polarity, chemotaxis, synapse formation, signaling, and effector responses; and hence it is not surprising that migration is a biomarker for cells with optimal effector functionality^32^. From a translational perspective, T cells from leukemia/lymphoma patients are known to harbor defects in migration and immunological synapse formation^33-35^. Since the FDA approved immunomodulatory drug, lenalidomide is known to rescue some of these defects, it will be important to quantify if addition of drugs like lenalidomide during CAR T cell manufacturing can yield fitter migratory cells with superior anti-tumor efficacy^34,36,37^.

T cells exhibiting migration have a high bioenergetic requirement to support locomotion. Our results confirmed that the migratory CAR T cells had enhanced mitochondrial SRC, a property that likely promotes long-term survival^38,39^. 3D confocal microscopy also demonstrated increased punctate mitochondria in individual cells with migration. Within this context, since AMPK is a well-known master regulator of SRC and mitochondrial remodeling, we confirmed that inhibiting AMPK has a profound impact on T-cell migration and consequently function at the single-cell level. To the best of our knowledge, this is the first direct demonstration of the role of AMPK in T-cell migration at the single-cell level, and this sets the stage for additional studies that explore this link between AMPK as a modulator of energetics and function within individual T cells and their antitumor efficacy. Our data is also consistent with studies that have demonstrated activated AMPK as a negative regulator of cellular adhesion and integrin activity in other cell types, but it is also important to recognize that AMPK is a global modulator of many different aspects of cell biology that need to be mapped within the context of T cells^40,41^.

An important question to consider in the context of IPs is how T-cell differentiation status impacts the migratory potential of T cells. Within our IP CD8^+^ T cells, scRNA-seq revealed that cluster CD8_6, enriched in CRs (**Figure 3**), has signatures of proliferation, migration and cytotoxicity. We compared these signatures to known T-cell differentiation signatures with the aid of STARTRAC (Single T-cell Analysis by Rna-seq and Tcr TRACking)^42^. STARTRAC performs paired scRNA-seq and TCR profiling (tracking clonotypes) to identify the properties of T cells that migrate and expand in the tumor relative to paired normal tissue/lymph nodes^42,43^. STARTRAC analyses of mouse and human colon/breast cancers have revealed that the migratory T cells are highly cytolytic and capable of expansion (similar to cells in cluster CD8_6), and are consistent with the TEM phenotype^42-44^. This needs to be interpreted with caution however, since as our sorting strategy with healthy donor-derived CAR T cells illustrates, migratory T cells are not confined to a single T-cell differentiation state. In comparing the signatures of CD4 and CD8 T cells within IP associated with clinical responses, a different hypothesis arises. While signatures of CD8 T cells are associated with immediate antitumor function including cytotoxicity, CD4 T cells signatures are primed for long-term persistence. Collectively, these data present a testable hypothesis of division of labor between the T cell subsets, and are consistent with emerging data for the role of CD4 CAR T cells in long term durable responses^21,45^.

Our results portray that migratory CAR T cells with increased bioactivity and functionality lead to persistence in vivo and control of tumor growth. There are, however, limitations to our study. We recognize that while the migration of these CAR^+^ T cells is a selectable property and reflective of the bioactivity of cells within a given population, it is unknown whether migration by itself can be utilized as a comparative marker in evaluating CAR^+^ T cell populations with varied CAR designs. Furthermore, while we have demonstrated the implementation of a simple transwell assay for the enrichment of migratory 19-28z T cells, as the data demonstrates, the segregation of cellular populations can be improved. For translational purposes, we anticipate building microfluidic chips capable of efficient segregation of large numbers of migratory T cells^46^. Lastly, while we recognize that migration might not necessarily be imprinted on T cells and their daughter cells, as they undergo cell division in vivo, migration nonetheless enables the identification and isolation segregation of T cells in vitro with long-lived potential in vivo.

In summary, we identified that migration is a marker that selects for T cells with superior antitumor effects reflective of productively activated cells with a balanced ability for functional execution without compromising proliferative potential. These attributes can be used to help improve manufacturing of T cells leading to predictable and superior patient clinical outcomes.

## METHODS

### Human subject statement

All work outlined in this report was performed according to protocols approved by the Institutional Review Boards at the University of Houston and the University of Texas MD. Anderson Cancer Center. TIL cells were expanded from surgical resection tissue under the protocol (2004–0069) approved by the Institutional Review Board (IRB) of the University of Texas MD Anderson Cancer Center (Houston, TX) and an FDA-approved Investigational New Drug (IND) application (NCT00338377).

### Patient samples

Patients with diffuse large B cell lymphoma (DLBCL) were treated with anti-CD19 CAR T cells. CAR T cells were made by Kite Pharma (Los Angeles, CA). After CAR T cell infusion, leftover cells in infusion bags were collected and transferred to the laboratory for microscopy assays and scRNA sequencing. At the 3-months follow-up, 10 patients showed complete response (CR, 62%), 2 showed partial response (PR, 13%) and 4 showed progressive disease (PD, 25%).

### Cell lines and primary T cells

Human pre-B cell leukemic line NALM-6 (ATCC) were cultured in T-cell medium (RPMI + 10% FBS) and used as CD19^+^ target cells. A second-generation CAR signaling via CD28 and CD3-ζ endodomains (with a mutated IgG4 spacer) were expressed in human T cells by electroporation with DNA plasmids from the *Sleeping Beauty* (SB) transposon/transposase system, as described previously^11^. T cells were used 2-3 weeks after transfection.

### Nanowell array fabrication

Nanowell array fabrication to investigate the effector functions at single-cell level was performed as described previously^47,48^. Briefly, we designed a master template of nanowell array using AutoCAD (Autodesk) and fabricated it on a silicon wafer using soft lithography techniques. We made the nanowell array by pouring Polydimethylphenylsiloxane (PDMS) on the silicon wafer and spinning the silicon wafer using a spin coater and then baking it at 70°C for 2 h. Then, we plasma oxidized the nanowell and attached it to the bottom of a 50-mm glass bottom Petri dish. Nanowell array had hundreds of nanowells, and each well was 50 µm × 50 µm × 50 µm in size.

### TIMING assays to profile functionality at single-cell level

We labeled approximately 1 million CAR T cells with PKH67 green fluorescent dye (Sigma) and 1 million NALM-6 cells with PKH26 red fluorescent dye (Sigma) as per the manufacturer’s protocol. We loaded both CAR T cells and NALM-6 cells on nanowell array in the way to reach mainly one effector and one to two targets per nanowell (E:T=1:1 or 1:2). We used complete cell-culture media (IMDM + 10% FBS) to cover the nanowell array. We added Annexin V-Alexa Fluor 647 (AF647) (Invitrogen(tm)) to the covering media to detect cell apoptosis during the experiment.

We used an inverted fluorescent microscope (Zeiss) which was equipped with a Lambda-DG4 illumination system, differential interference contrast (DIC) condenser annulus, 20× 0.80 NA Zeiss Plan-Apochromat objective and Orca Flash 4.0 camera (Hamamatsu). We placed the petri dish containing the PDMS-nanowell array, on a motorized stage in a box to maintain the temperature at 37°C and CO2 level at 5%. We used Alexa Flour (488 nm), TexasRed (566 nm) and Cy5 (651 nm) channels for detection of CAR T cells, NALM-6 cells, and apoptosis, respectively. Images were acquired for six hours with five-minute intervals.

### Confocal microscopy for integrated measurements of subcellular organelles

For confocal microscopy, we fluorescently labeled ∼300,000 CAR T cells at 37°C for 30 minutes in 1:1 (v/v) solution of live cell staining buffer (Abcam) and RPMI-1640 (Corning) containing final concentrations of 500 nM MitoTracker(tm) Deep Red FM (Invitrogen(tm)), 250 nM LysoRed (Abcam) and 1 µM Hoechst 33342 (Sigma) for labeling mitochondria, lysosome, and nucleus, respectively. Then we washed cells with Hanks’ balanced salt solution (Cellgro) + 10% HEPES (Corning) and resuspended in cell culture media (RPMI + 10% FBS), and then loaded labeled cells on a glass-bottom 96 well plate (MatTek Corporation) for confocal imaging.

We used a Nikon A1/TiE inverted microscope equipped with a 100x, 1.45 NA objective for imaging. We took 3D images (∼50 slices, 0.3 µm/slice) from multiple fields of view using DAPI (404.0 nm), FITC (488.0 nm), TexasRed (561.8 nm) and Cy5 (641.0 nm) channels for detection of nucleus, membrane, lysosome, and mitochondria, respectively.

### SImage processing, cell segmentation, cell tracking, and data analytics

For TIMING assay image processing, we used a home-made pipeline to analyze 16-bit images as described before^49^. Briefly, an automated pipeline was implemented for automatic detection of nanowells, cell segmentation, tracking and feature computation. The pipeline output are the tables containing statistical information for nanowells with one effector cell and up to five target cells (1E:1T – 1E:5T). We partitioned events based on the functionalities of the cells i.e. 1E:1T-killing, mono-killing and serial killing (**Figure S2A-B**):

a. 1E:1T killing: a single T cell killing a target cell existing in the nanowell upon conjugation.
b. Mono killing: a single T cell killing only one target cell upon conjugation, while multiple targets (at least two) exist in the nanowell.
c. Serial killing: a single T cell killing at least two target cells upon conjugation, while multiple targets (at least two) exist in the nanowell.
d. No killing: T cells without killing any target cells despite evidence of conjugation.

For tracking the mitochondria and lysosomes within the cells using confocal microscopy, Z-stacks of 16-bit images were extracted for each channel and processed in ImageJ (National Institutes of Health) using a series of plugins. First, the Subtract Background plugin was applied to the mitochondria and lysosome channels prior to segmentation to reduce variations in background intensities. Next, the 3D Objects Counter plugin was applied to the background-corrected image to determine mitochondrial and lysosome regions of interest (ROIs). ROIs were overlaid onto the original image and measurements were collected. Similarly, the 3D Objects Counter plugin was used on nucleus channel using the original image only. Lastly, tracking of single cell movement was done using the TrackMate plugin^50^ to filter out unstable cells upon their movement. All measurements were consolidated in R, where mitochondria and nuclei were matched to their corresponding cell.

### Single cell RNA sequencing (scRNA-seq)

We performed scRNA-seq for nine samples (5 CR and 4 PD) with enough number of residual cells. We first used dead cell removal kit (Miltenyi Biotec, Germany) containing MicroBeads for the magnetic labeling of dead cells and we then removed the dead cells by passing the resuspended cells through the magnetic field of a MACS separator. We performed library preparation process in three batches. For each batch, we used three different TotalSeq C anti-human Hashtag Antibodies (BioLegend, San Diego, CA) to multiplex the samples as per the manufacturer’s protocols. Next, we did transcriptome and TCR capturing using 10x Chromium platform (10x Genomics, Pleasanton, CA). We used Chromium single cell 5’ reagent v2 kit for gene expression and V(D)J profiling. The sequencing was performed using a HiSeq PE150 sequencer (Illumina, San Diego, CA).

### ScRNA-seq analysis

We processed gene expression FASTQ files generated with Illumina sequencer using the Cell Ranger pipeline (version 6.0.0, 10x Genomics) for read alignment and generation of feature-barcode matrices. The output files then were uploaded into R (version 4.0.1) for further processing using Seurat Package (version 4.1.0)^51^. We used the SAVER package^52^ first, to recover the gene expression profile in the data. Then we filtered out the cells with high mitochondrial gene expression (more than 15% of the read counts) and we ended up with 21,469 cells from nine patients with mean unique molecular identifier (UMI) of 7738.

We detected the highly variable genes and significant principal components (PCs) following the Seurat standard workflow for unsupervised clustering of the cells and used uniform manifold approximation and projection (UMAP) for visualization of the clusters. We identified CD8 and CD4 T cell subsets using *CD8A, CD8B, CD4* and *CD40LG* gene markers. We identified CAR^+^ cells by detection of the CAR sequence (FMC63-CD19scFV, GenBank: HM852952.1).

We used gene set variation analysis (GSVA) package^53^ in R to calculate ssGSEA scores for different pathways. We used pre-defined gene-sets from molecular signatures database (MSigDB, v7.5.1) for pathway analysis. We also used customized gene-sets: TCF7.regulon^17^, T cell migration^18^, CD4^+^ Ki67^21^ and CD8^+^ effector^42^ based on previous publications. Lowly expressed genes (average expression <0.25) were removed from the pathway analysis. We generated heatmaps by pheatmap package in R. We performed statistical analysis and generated *P* values in R.

### Migration of T cells through a transwell migration chamber

Unstimulated, overnight serum deprived, CAR^+^ T cells were seeded on the top compartment of PET five or eight μm-pore Boyden transmigration chamber (EMD Millipore), while the lower compartment contained FBS rich media. After 4-6 hours, the cells from the bottom and the top compartment were harvested as “migratory” and “non-migratory” populations, respectively. The lower part of the membrane was washed into the “migratory” cell suspension, while the top surface of the membrane is washed into the “non-migratory” cell suspension. We analyzed the phenotype and function of the cells using flow cytometry and TIMING.

### Flow cytometry-based phenotyping

For phenotyping, we stained the CAR^+^ T cells using a panel of human-specific antibodies CD62L (DREG-56), CD45RA (HI100), CD3 (SK7). CD4 (OKT4), CD8 (RPA-T8) and Granzyme B (QA16A02) from Biolegend. The anti-CAR scFv was made in house^54^. We analyzed the cells using BD LSRFortessa X-20 cell analyzer.

### Compound C inhibition assays

We incubated T cells with 10 μM dorsomorphin (Sigma Aldrich) for a period of 6-24 hours. The T cells were subsequently used for either migration assays or functional profiling using TIMING assays. Incubation with compound C did not have an impact on T-cell viability (data not shown).

### In vivo efficacy of CAR^+^ T cells

On day 0, 7-week-old NOD.Cg-PrkdcscidIl2rgtm1wjl/SzJ (NSG) mice were injected intravenously (i.v.) via a tail vein with 1.5×10^4^ EGFP^+^ ffLuc^+^ NALM-6 cells. Mice (n = 10/group) in the two treatment cohorts received via tail vein injection (on day 5) of 10^7^ CAR^+^ T cells. One group of mice (n = 10) bearing tumor were not treated with T cells. Anesthetized mice underwent bioluminescent imaging (BLI) in an anterior-posterior position using a Xenogen IVIS 100 series system (Caliper Life Sciences) 10 minutes after subcutaneous injection (at neck and shoulder) of 150 μL (200 μg/mouse) freshly thawed aqueous solution of d-Luciferin potassium salt (Caliper Life Sciences) as previously described^55^. Photons emitted from NALM-6 xenografts were serially quantified using the Living Image 2.50.1 (Caliper Life Sciences) program. On day 28, five mice in each group were euthanized to evaluate the presence of T cells and tumor cells. Bone marrow was flushed from the femurs using 30Gx1/2inch needles (BD, catalog no. 305106) with 2% FBS in PBS. Spleens were disrupted using a syringe in 2% FBS/PBS and passed through a 40 μm nylon cell strainer (BD, catalog no. 352340) to obtain a single-cell suspension. Red blood cells from bone marrow, spleen, and peripheral blood were lysed using ACK lysing buffer (Gibco-Invitrogen, A10492) and remaining cells were stained for the presence of tumor (human CD19 and EGFP), T cells (human CD3) and CAR T cells (scFv) by flow cytometry. The remaining five mice in each group were used to determine the survival curves. In the suboptimal dose model, the mice were treated exactly as above except that on day 5, 2×10^6^ CAR^+^ T cells were injected intravenously.

### Data visualization and statistical analysis

Data plotting and statistical analysis were performed in R and GraphPad Prism v7. Schematics were made in Inkscape (v1.1.2).

## AUTHORSHIP CONTRIBUTION

Designed the study: AR, GR, SN, LJNC, HS and NV

Prepared the manuscript: AR, GR, MF, HS, SN, NV and LJNC

Performed experiments: AR, GR, MF, MMP, KF, XA, FS and IB

Analyzed data: AR, GR, MF, AS, MMP, XA, NA and FS

Provided patient samples: HS, LJNC, SN, NPO, AB, CB, MM and DH

All authors edited and approved the manuscript.

## ACKNOWLEDGEMENTS

This publication was supported by the NIH (R01GM143243); CPRIT (RP180466); MRA Established Investigator Award to NV (509800), NSF (1705464); CDMRP (CA160591); and Owens foundation. We would like to acknowledge the MDACC Flow Cytometry and Cellular Imaging Core facility for the FACS sorting (NCI P30CA16672), Intel for the loan of computing cluster, and the UH Center for Advanced Computing and Data Systems (CACDS) for high-performance computing facilities.

