## Supplementary information for "T-cell potential for CD19-expressing malignancies revealed by multi-dimensional single-cell profiling"

**Table ST1. Glossary of terms used in the paper**

| | |
|---|---|
| CAR | Chimeric Antigen Receptor |
| 19-28z | CD19-specific CAR construct with an IgG4 spacer and CD28 and CD3-ζ endodomains |
| Migration | Persistent motion wherein the direction of movement is maintained for at least one cell diameter |
| LBCL | Large B-cell lymphoma |
| DLBCL | Diffuse large B-cell lymphoma |
| Axi-cel | Axicabtagene ciloleucel |
| E | Effector $CAR^+$ T-cell |
| T | Target cell |
| TIMING | Timelapse Imaging Microscopy In Nanowell Grids |
| Non-killer | T cells that do not kill any tumor cells despite evidence of conjugation |
| Mono-killing | Ability of a single T-cell to kill exactly one target cells |
| Multi-killing | Ability of a single T-cell to kill two or more target cells |
| Mono-killer | T cells that kill exactly one tumor cell at an E:T ratio of 1:2-5 |
| Serial killer | T cells that kill at least two tumor cells at an E:T ratio of 1:2-5 |
| Killing efficiency | Description of the kinetics of killing mediated by individual T cells (Please see $t_{Death}$ below) |
| Conjugation | Stable contact between effector cell and target cell lasting > 5 minutes |
| $t_{Seek}$ | Time taken for the effector cell to conjugate to tumor cell |
| $t_{Death}$ | Time elapsed between first conjugation and tumor cell apoptosis (Annexin V staining positive) |
| $t_{Conjugation}$ | Cumulative duration of conjugation between $t_{Seek}$ and $t_{Death}$ |
| AR | Aspect ratio of polarization represented as the ratio of the minor and major axes of the cell, fitted to an ellipse |

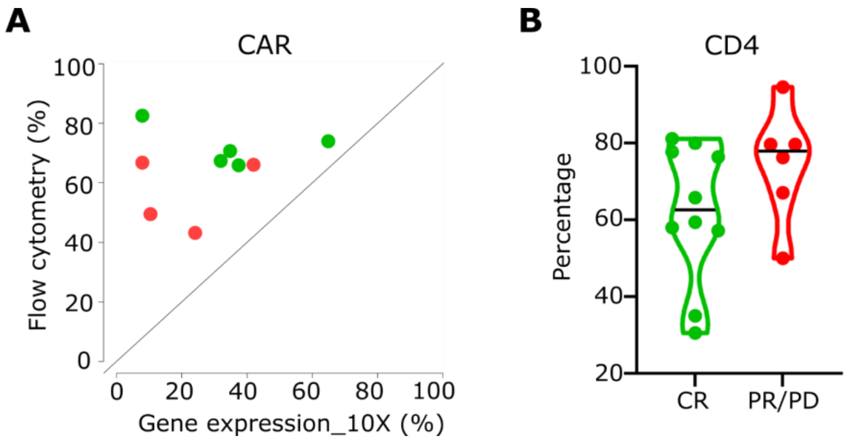


**Figure S1. Phenotype characteristics measured by flow cytometry and scRNA-seq.**

(A) Comparisons of $CAR^+$ T cells identified by flow cytometry and mRNA expression for nine patients.

(B) Comparisons of $CD4^+$ T cells recorded by flow cytometry for all sixteen patients. There is no significant difference between CR and PR/PD.

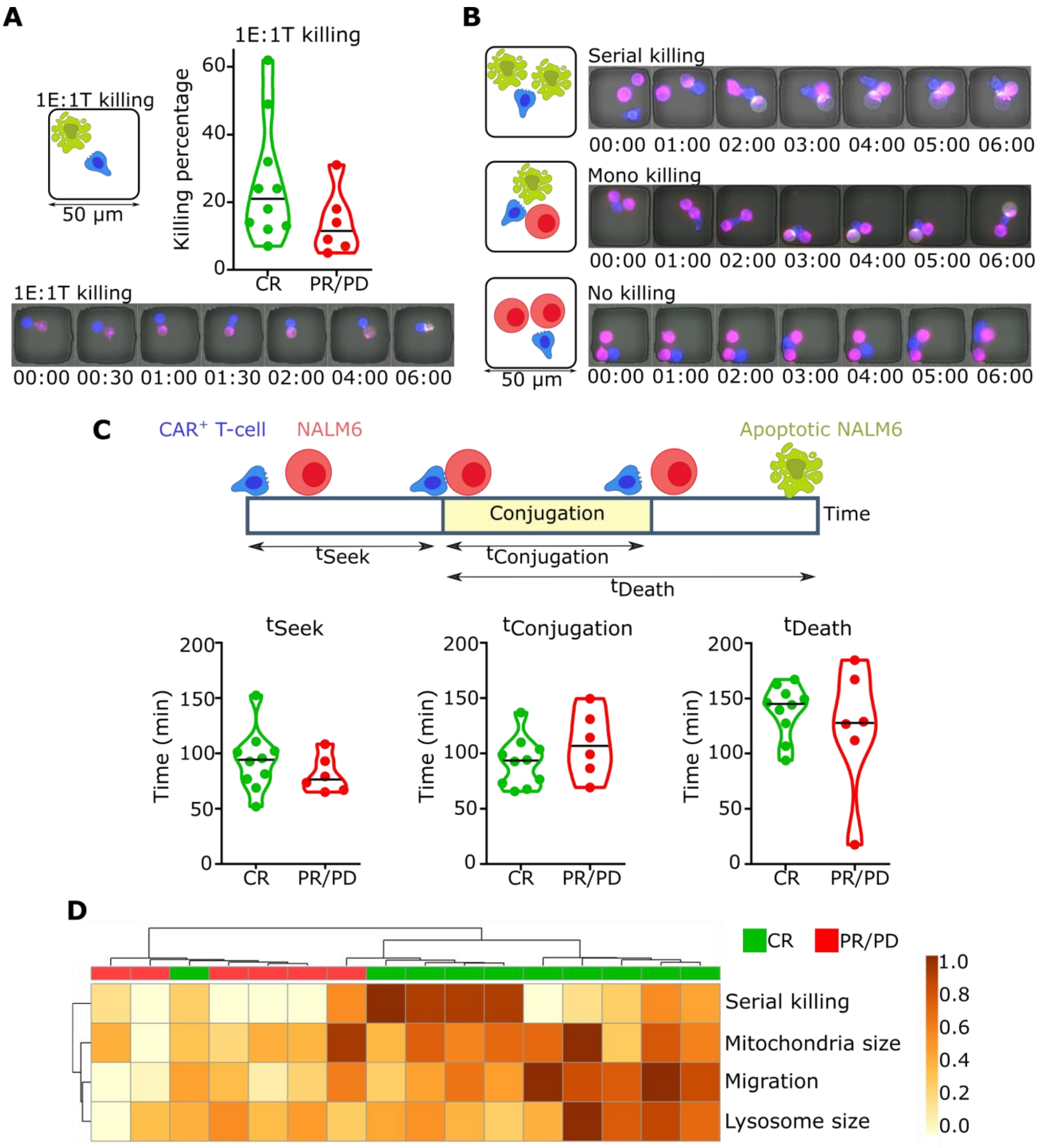


**Figure S2. T cells from CR are enriched for serial killing, lysosomal size and persistent migration.**

(A) Schematic of a killing event at an E:T of 1:1 in which a CAR T cell conjugates and kills a NALM-6 cell. The plot on the right shows the killing rate comparison between T cells from CR and PR/PD within all 1E:1T nanowells. Micrograph showing an example of 1E:1T killing event through the 6-hours (hh:mm) of time-lapse imaging.

(B) Schematics and examples of serial killing, mono killing and no-killing events (all possible outcomes in the presence of conjugation) in nanowells with an E:T of 1:2.

(C) Schematic of a killing event, showing the interaction parameters. $t_{Seek}$ defined as the time for CAR T cell to find and conjugate to the NALM-6 cell. $t_{conjugation}$ is defined as the duration of CAR T cell in stable conjugation with NALM-6 cell. $t_{Death}$ is the time interval between the start of the conjugation and the apoptosis of the NALM-6 cell. Plots show the comparison between T cells from CR and PD/PR for these parameters. There was no significant difference between T cells from CR and PR/PD in terms of dynamics of interaction.

(D) Unsupervised hierarchical clustering based on parameters from TIMING, and confocal microscopy. Serial killing, migration, mitochondrial volume and lysosomal volume were features associated with T cells from CR patients.

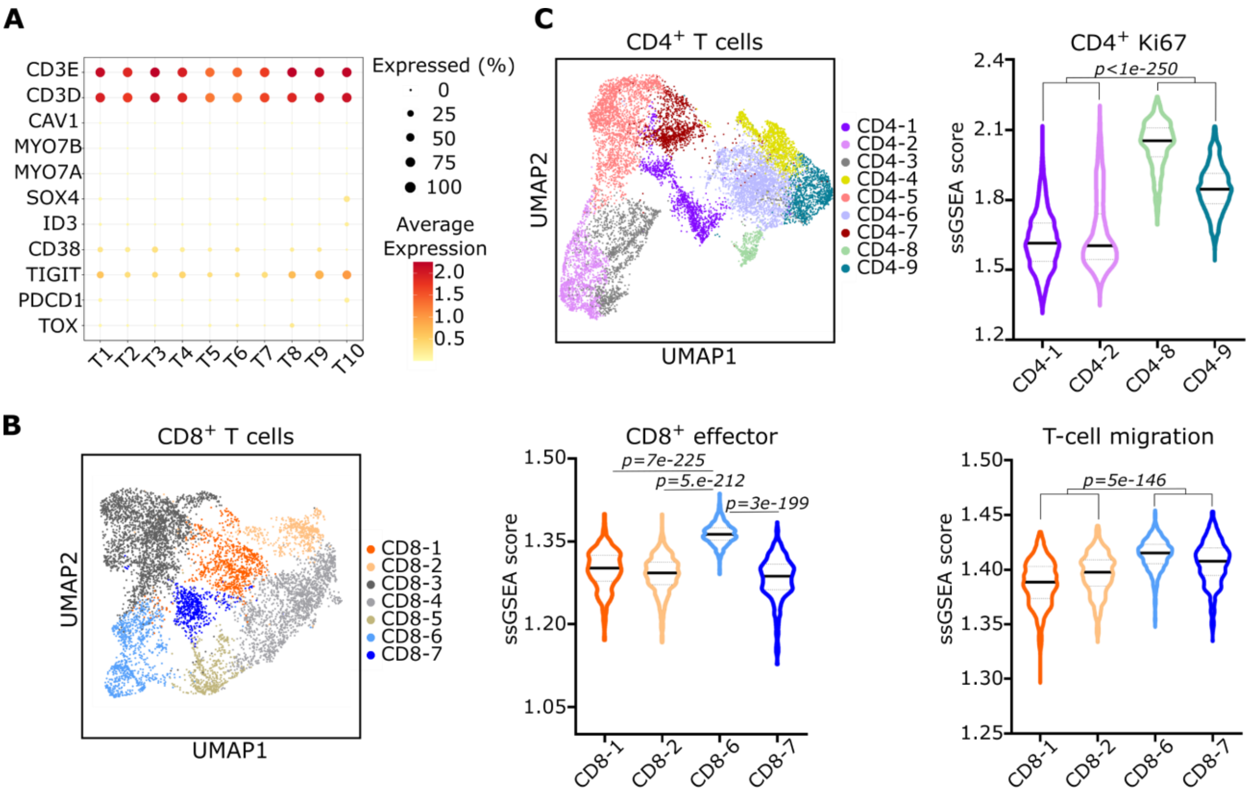


**Figure S3. T cell phenotypes defined using scRNA-seq.**

(A) Bubble plot showing key genes associated with T cell migration and exhaustion phenotypes. Independent of clinical response, these genes did not show significant expression within any of the clusters.

(B) *UMAP for CD8$^{+}$ T cells (n=7,439)*. Seven clusters identified using unsupervised clustering. Violin plots showing the ssGSEA score for migration and CD8 effector phenotypes calculated for two clusters enriched in PD (CD8-1 and CD8-2) and two clusters enriched in CR (CD8-6 and CD8-7). T cells from the CD8-6 cluster show higher effector signatures than the other three clusters. For migration, CD8-6 and CD8-7 show significantly higher signatures compared to the CD8-1 and CD8-2.

(C) *UMAP for CD4$^{+}$ T cells (n=12,527).* Nine clusters identified using unsupervised clustering. Violin plot showing the ssGSEA score for long-persisting Ki67$^{high}$ CD4$^{+}$ CAR T phenotype calculated for two clusters enriched in PD (CD4-1 and CD4-2) and two clusters enriched in CR (CD4-8 and CD4-9). T cells from CD4-8 and CD4-9 clusters show significantly higher long-persistence signatures compared to the CD4-1 and CD4-2 clusters.

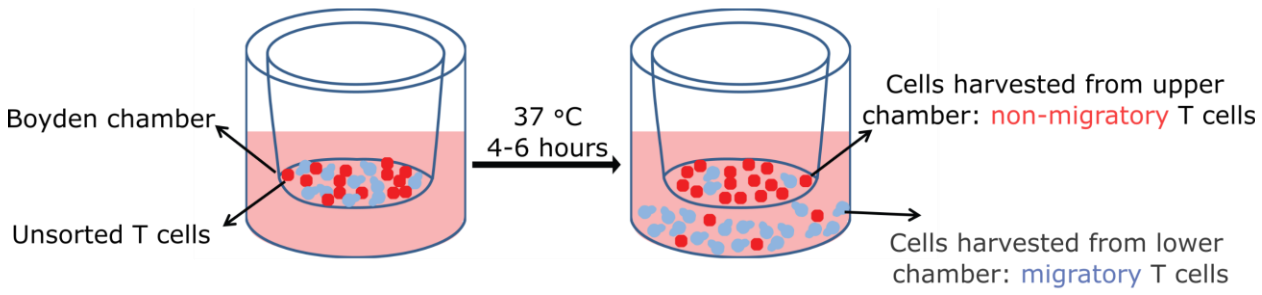


**Figure S4.** Modified transwell assay for the isolation of migratory 19-28z T cells. 19-28z T cells are seeded onto the top well of a transwell chamber. After 4-6 hours, the migratory 19-28z T cells are harvested from the bottom and the non-migratory 19-28z T cells are retrieved from the top of the chamber. Immediately after this segregation, the cells are used in functional experiments or infused in vivo with no further culturing/alterations.

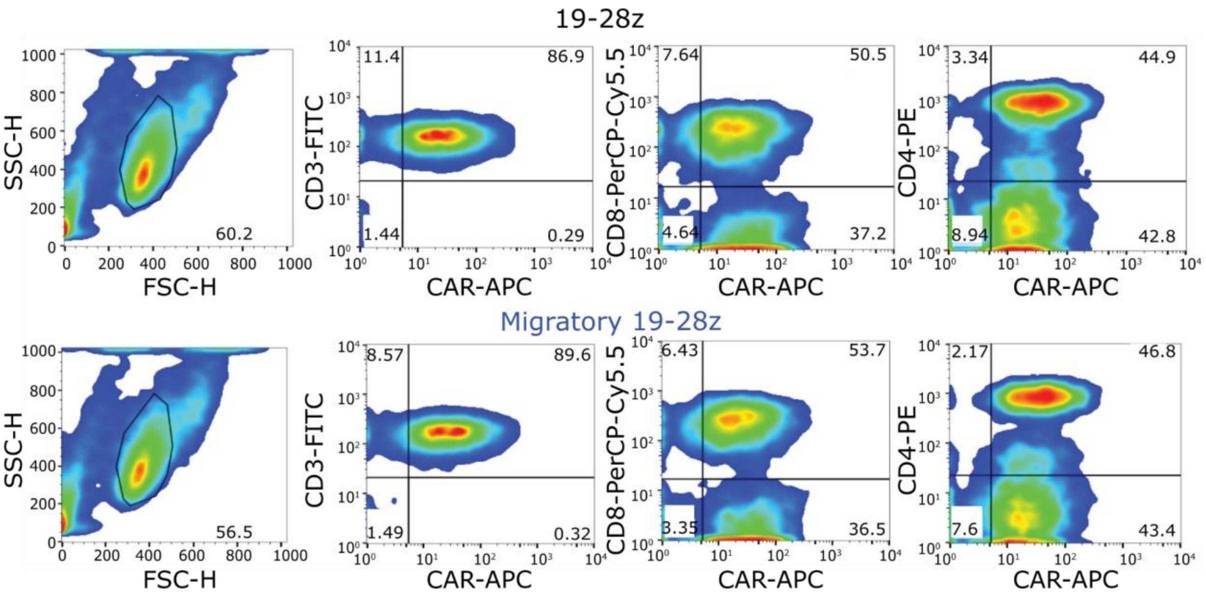


**Figure S5.** A representative example of the phenotype of the matched 19-28z and migratory 19-28z T cell populations. This data is representative of at least four healthy donor-derived T cell populations.

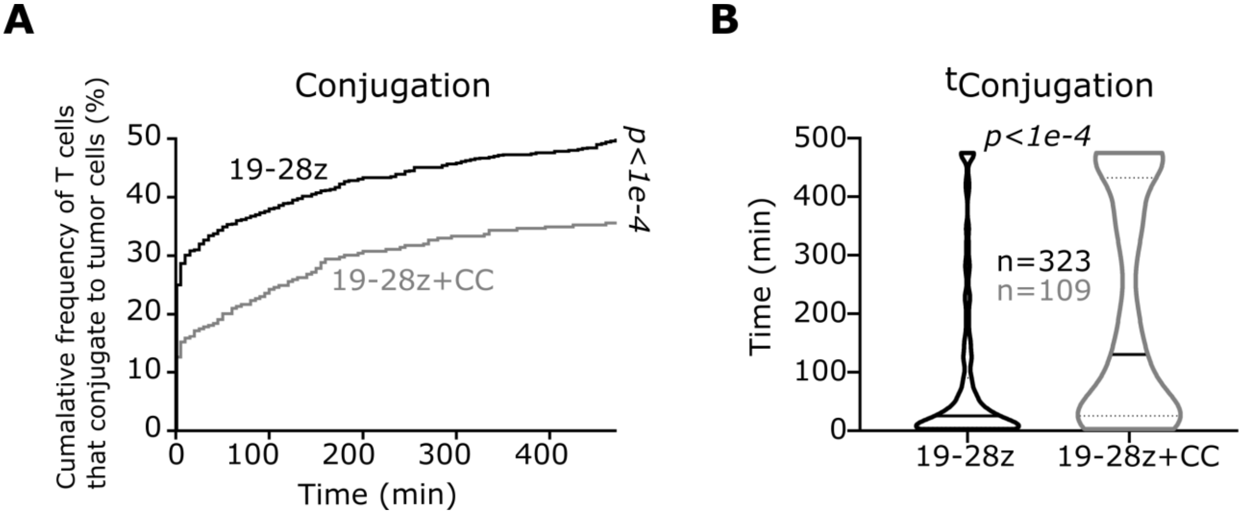


**Figure S6. The impact of AMPK inhibition on 19-28z T cell antitumor function revealed by TIMING.** Upon CC treatment, 19-28z T cells displayed:

(A) Lower frequencies of conjugation to tumor cells

(B) Extended duration of contact subsequent to conjugation of tumor cells in comparison to control (DMSO) 19-28z T cells. Each data point represents a single-cell. The black bar represents the median and the dotted lines denote quartiles in B. *P* values in A and B were computed using log-rank tests and a two-tailed t-test, respectively.

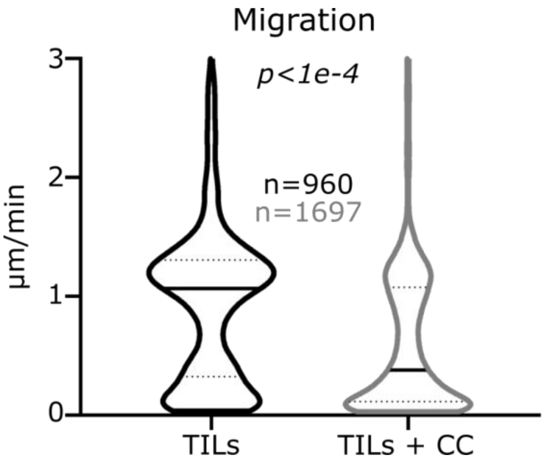


**Figure S7. The impact of AMPK inhibition on melanoma-derived, expanded tumor infiltrating lymphocytes (TILs).** Upon CC treatment TILs demonstrated a significant reduction in directional migration. Each data point represents average migration for a single cell. The black bar represents the median and the dotted lines denote quartiles. *P* value was computed using Mann-Whitney tests.

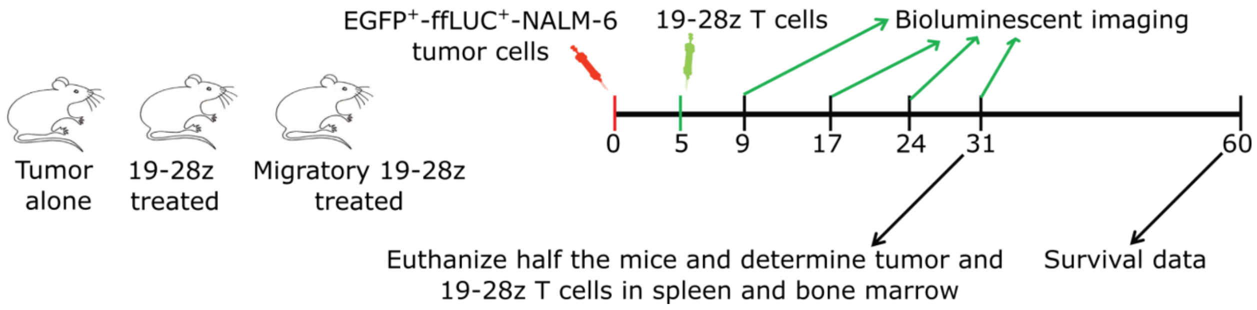


**Figure S8. Design of mice experiments to determine the relative efficacy of the 19-28z populations.** Groups of ten mice were treated with either 19-28z T cells or migratory 19-28z T cells five days after the injection of ffLuc expressing EGFP$^{+}$NALM-6 cells. One set of five mice from each of the two treated groups were sacrificed at day 31 to quantify both tumor cells and persisting T cells within the spleen and bone marrow of the mice.

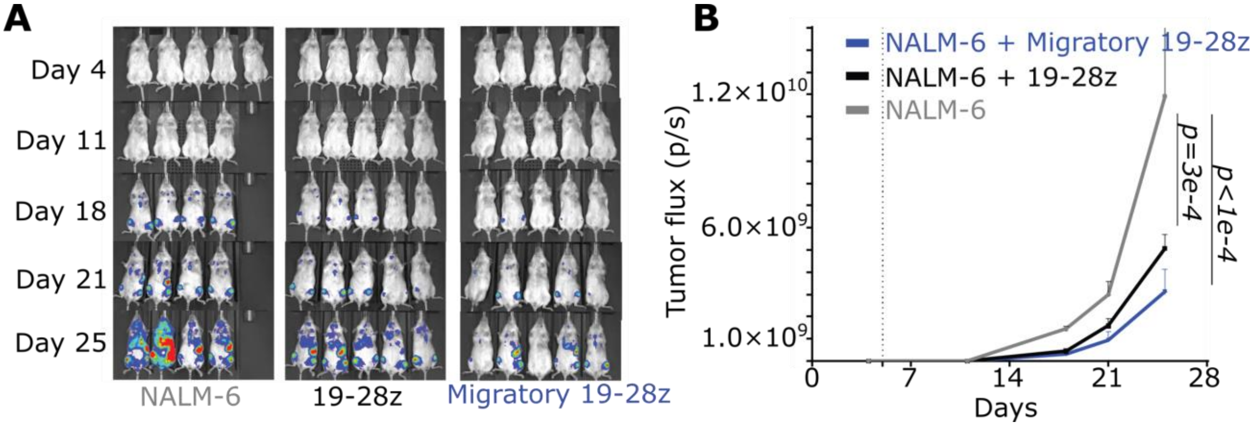


**Figure S9. Migratory 19-28z T cells showed enhanced antitumor activity compared to the unsorted 19-28z T cells in suboptimal dose model.**

(A) False-colored images illustrating the photon flux from ffLuc expressing $EGFP^{+}$NALM-6 cells treated with suboptimal doses of 19-28z T cells.

(B) Time course of the longitudinal measurements of NALM-6 derived photon flux from the three separate cohorts of mice (n= 5 in each group). The background luminescence was defined based on mice with no tumor. Error bars represent SEM and *P* values were computed using a two-way ANOVA.

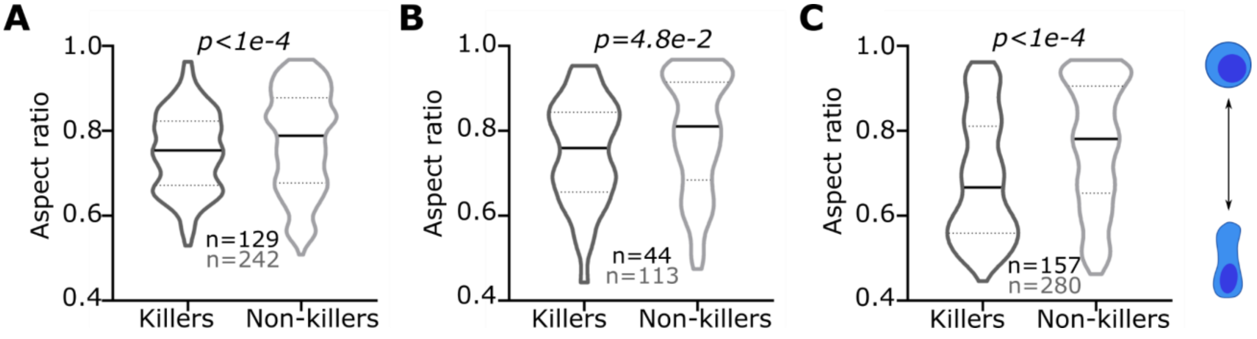


**Figure S10. Quantifying the link between polarization and functionality in diverse CARs.**

(A, B and C) The polarization of individual killer and non-killer CAR T cells without and with conjugation to tumor cells. All data from an E:T of 1:1. (A) shows the data for 19-8-28z T cells tested against NALM-6 cells. (B) and (C) show the data for two different constructs of tri-specific CAR$^+$ T cells tested against patient-derived tumor cells. The black bar represents the median and the dotted lines denote quartiles. All *P* values were computed using Mann-Whitney tests and each data point represents a single effector cell.
